# Nuclear Lipid Droplets Exit to the Cytoplasm: A Novel Mechanism of non-nuclear pore-dependent material exchange

**DOI:** 10.1101/2023.09.08.556761

**Authors:** Yi Jin, Jin Huang, Zichen Zhao, Yanjie Tan, Jian Wu, Zhuqing Ren

## Abstract

In cellular biology, the extrusion of large molecular complexes from the nucleus has consistently represented a considerable challenge. In this investigation, we reveal a hitherto unexplored mechanism facilitating the efflux of nuclear lipid droplets (nLDs) from the nucleus directly into the cytoplasm, a process integral to comprehending cellular processes and nuclear dynamics. Through high-throughput sequencing, immunoprecipitation, and immunofluorescence experimentation, we demonstrated that the type I nucleoplasmic reticulum engages with nLDs, enabling the dynamic protein Rab8a’s transfer to the nLDs and fostering their translocation to the nuclear membrane. Phosphorylation of LMNA/C at the S392 site, adjacent to the nLD-nuclear membrane contact site, induces local depolymerization of lamins, creating an aperture that enables gradual nuclear exit of nLDs. Following this, nLDs, carrying DNA fragments, are released into the cytoplasm through budding and subsequently undergo autophagy-mediated degradation. Furthermore, we demonstrate that RhoA-PKC signaling enhances nLD movement and LMNA/C phosphorylation, while its inhibition impedes nuclear nLD efflux. Our groundbreaking investigation illuminates non-nuclear pore-dependent mechanisms of material exchange and offers fresh perspectives on nuclear metabolic pathways.

**In Brief:** 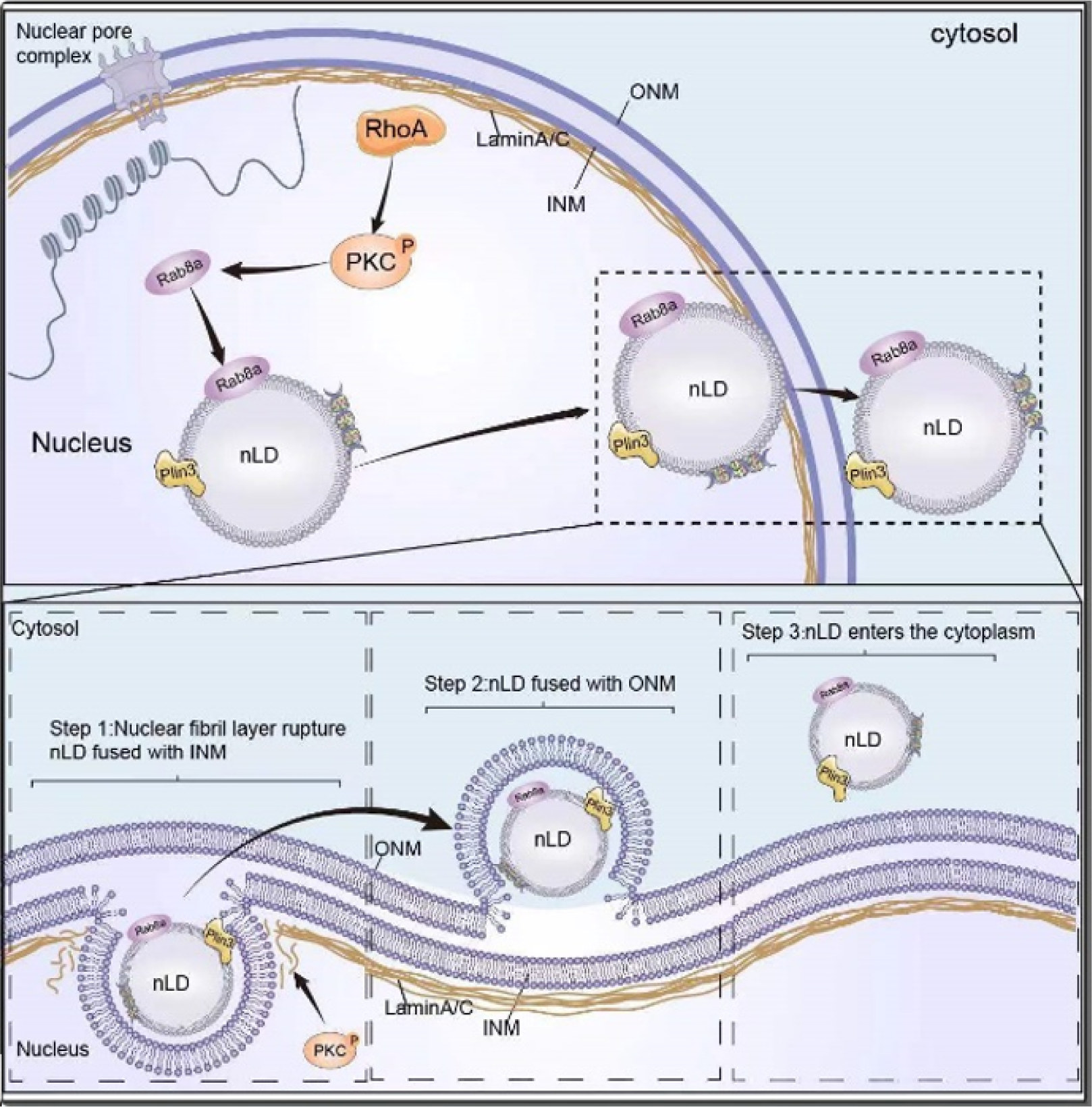

In this study, Jin et al. elucidate a novel mechanism for the expulsion of nuclear lipid droplets (nLDs) from the nucleus. They demonstrate that the activation of RhoA within the nucleus recruits Rab8a to the surface of nLDs. Concurrently, PKC phosphorylation triggers the phosphorylation of LMNA/C at the S392 site, leading to a localized depolymerization of lamins. This creates an aperture that allows for the gradual exit of nLDs from the nucleus. The efflux process of nLDs involves three sequential steps: rupture of the nuclear fibril layer, fusion of nLDs with the inner nuclear membrane (INM), and subsequent fusion with the outer nuclear membrane (ONM). This enables nLDs to enter the cytoplasm, shedding light on a previously unknown process of large molecular entity expulsion from the nucleus.

## Introduction

The cell nucleus is not only a central repository for the storage and transmission of genetic information but also acts as a site for key metabolic processes. Central to the nucleus’s function is the regulation of its contents, which is maintained by rigorously controlled mechanisms of substance transport. These mechanisms enable the movement of proteins, RNAs, and a diverse array of molecules, including lipids, across the nuclear envelope. They rely on nuclear pore complexes for active transport and use passive diffusion or vesicular transport for smaller, non-polar, and uncharged substances (Akey et al., 2022; Fernandez-Martinez et al., 2016; Fontana et al., 2022; Huang et al., 2022; Manik et al., 2017; Zhu et al., 2022).

In recent years, lipid metabolism has gained prominence, especially regarding the roles and dynamics of lipid droplets (LDs). These multifunctional organelles, known for their diverse cellular functions—energy storage, protein maturation, lipid precursor synthesis, and cellular stress regulation (Olzmann and Carvalho, 2019; Zhang and Liu, 2017)—have traditionally been linked with the cytoplasm. However, recent discoveries have revealed the existence of nuclear lipid droplets (nLDs) within the nucleus under specific physiological and pathological conditions (Romanauska and Kohler, 2018, 2021; Soltysik et al., 2019; Soltysik et al., 2021).

The formation of nLDs takes place through lipid metabolism within the inner nuclear membrane, where lipid metabolites are synthesized into glycerol esters and released into the nucleoplasm, resulting in nLD genesis (Romanauska and Kohler, 2018). These organelles play a vital role in maintaining lipid homeostasis in the nucleus and alleviating nuclear lipid metabolism stress. For instance, previous studies have indicated that CCT-α, a protein involved in phosphatidylcholine synthesis, is recruited to the surface of nLDs, contributing to the mitigation of endoplasmic reticulum stress (Soltysik et al., 2019). Moreover, conditions of elevated unsaturated fatty acids or impaired cellular phosphatidylcholine synthesis have led to the emergence of nLDs (Romanauska and Kohler, 2018, 2021; Soltysik et al., 2019; Soltysik et al., 2021). The importance of these dynamics is emphasized by observations of abnormal nLD formation in pathological conditions, such as nonalcoholic fatty liver disease (NAFLD), a widespread and often debilitating condition that links disrupted cellular lipid metabolism with nLD formation (Ostlund et al., 2022; Shin et al., 2019).

In light of this context, understanding the metabolic mechanisms and regulatory roles of nLDs in cellular lipid metabolism emerges as a critical endeavor. Our study, therefore, unveils novel findings on the process of nLD egress from the nucleus. Intriguingly, we observed that nLDs do not disassemble within the nucleus but are instead directly expelled, a process typically associated with the transport of smaller nucleolar materials through nuclear pores. This astonishing discovery of the direct exit of larger organelles, such as LDs (500 nm-1 µm), from the nucleus, enriches our understanding of the intricate relationship between nuclear metabolism and material exchange. By unraveling this previously unknown mechanism of LD exit, our study provides invaluable insights into the complex interplay between nuclear metabolism and material exchange, which could potentially advance our understanding of diseases related to aberrant lipid metabolism.

## Materials and Methods

### Cell culture and transfection

The Huh7 cell line was purchased from Type Culture Collection of the Chinese Academy of Sciences (Wuhan, China). Huh7 cells were cultured in Dulbecco’s modified Eagle’s medium (DMEM; HyClone, Logan, UT, USA) with 10% fetal bovine serum (FBS; #SH30396.03, Hyclone, Canada), 100 unit/mL penicillin, and 100 μg/mL streptomycin in dishes at 37 °C in a humidified atmosphere with 5% CO2.

For transfection, cells were seeded in a 6-well plate or on slides in a 24-well plate. Then, the cells were transfected with Lipo2000™ Transfection Reagent (Invitrogen, Shanghai, China). For the preparation of RNAi working solution, 10 μL siRNA oligo (20 μM, Ribobio, Guangzhou, China) was mixed with 10 μL Lipo2000 reagent in 100 μL Opti MEM. For the preparation of the overexpression of the working solution, 2.5 μg plasmid was mixed with 4 μL Lipo2000 reagent in 50 μL Opti MEM. The working solution was added to the plate’s wells and incubated for 6 h. Then, the plate’s wells were changed with fresh cultural medium (DMEM with 10% FBS) for another 48 h of culture.

### Protein mass spectrometry

The protein mass spectrometry detection for this study was commissioned by Novogene Technology Co., Ltd., Beijing, China. In brief, protein alkylation reduction and enzymatic hydrolysis were first performed, followed by peptide desalting, and mass spectrometric detection. The obtained mass spectrometry data were subjected to further quality analysis and protein function annotation for further cluster analysis. For testing methods and specific parameter settings, we can provide detailed instructions if necessary.

### Nuclear LDs and cytosol LDs isolation

For cytosol LD isolation, we first used the cytoplasmic separation kit to separate the cytoplasmic group, and then we isolated and purified LDs in the cytoplasm according to Ding’s method (Ding et al., 2013). Briefly, we added the cytoplasmic components to Buffer A, and then ultracentrifuged (185,000× g) for 2 hours and took the upper LDs. The LD components obtained after ultracentrifugation were washed 3 times with Buffer B, each time using 15,000× g centrifugation, and, finally, the supernatant was discarded. The obtained LD fractions were used to extract proteins using a protein lysate.

For nLD isolation, the whole nuclei were isolated using a kit (#SN0020, Solarbio, Beijing, China). Then the nuclei were washed three times with PBS buffer. Thereafter, the integrity of the nuclei and the purity were detected by fluorescence microscope. The nuclei were treated with 0.8% Triton X-100 for 15 min, then grinded on ice 50 times in Buffer A. The nLDs were isolated through ultracentrifugation (185,000× g) for 2 hours and took the upper LDs. The LD components obtained after ultracentrifugation were washed 3 times with Buffer B, each time using 15,000× g centrifugation, and, finally, the supernatant was discarded. The obtained LD fractions were used to extract proteins using a protein lysate.

### Western blot

Western blotting was performed as reported previously (Karthika et al., 2016; Tan et al., 2019). Briefly, cells were collected and homogenized in lysis buffer (#P0013, Beyotime Biotechnology, Nanjing, China). Then, the homogenates were incubated with an SDS-PAGE sample loading buffer (#P0015A, Beyotime Biotechnology, Nanjing, China) at 98 °C for 10 min. Subsequently, the samples were separated by 10% sodium dodecyl sulfate-polyacrylamide gel electrophoresis (SDS-PAGE) and were transferred to a polyvinylidene fluoride (PVDF) membrane (Bio-Rad, USA) using a semidry electrophoretic apparatus. The blocked membranes (#P0252-100mL, QuickBlock™ Blocking Buffer for Western Blot, Beyotime Biotechnology, Nanjing, China) were incubated with antibodies overnight at 4 °C. The blots were extensively washed three times with tris-buffered saline with Tween 20 (TBST) buffer for 10 min and were incubated under gentle agitation with the primary antibodies for immunodetection at 37 °C for 1.5 h (diluted in QuickBlock™ Primary Antibody Dilution Buffer for Western Blot, #P0256, Beyotime Biotechnology, Nanjing, China). Then, the blots were extensively washed three times with TBST. Subsequently, blots were incubated under gentle agitation with the secondary antibodies for immunodetection at 37 °C for 1 h (diluted in QuickBlock™ Secondary Antibody Dilution Buffer for Western Blot, #P0258, Beyotime Biotechnology, Nanjing, China). For detection, an M5 ECL Western Blot Kit (#MF-078-01, Mei5bio, Beijing, China) and the chemiluminescence imaging system (LAS4000, ImageQuant, Germany) were used. The original western blots are showed in Supplemental Material.

### LDs marking

LDs marking was performed as reported previously (Tan et al., 2019). The cell slides were fixed with 4% paraformaldehyde for 15 min at room temperature. The slides were stained with BODIPY 493/503 (#D3922, Invitrogen, Carlsbad, CA, USA) for 10 min at 37 °C and were then stained with DAPI for 10 min at 37 °C. After washing three times with PBS for 10 min each, the slides were sealed with an anti-fluorescent quenching solution (#P36961, ProLong™ Diamond Antifade Mountant, Invitrogen, Thermo Fisher, USA) for confocal microscopic observation (63× oil lens, BODIPY FL and DAPI channels, Zeiss LSM 800, Germany).

### Oleic acid medium treatment

Oleic acid treatment was as described in previous studies (Tan et al., 2019). For the oleic acid treatment, a 20 mM oleic acid–phosphate-buffered saline (PBS) mixture and 20% FA-free bovine serum albumin (BSA) medium were prepared, and both media were heated in a 70 °C water bath for 30 min. Finally, the media were mixed. The 10 mM oleic acid–BSA mixture was added to the cell cultural medium at 1:49 (v:v). The cells were then either seeded on slides or on plates that had been washed three times using PBS. Then, 1 mL oleic acid medium was added to the well, and the cells were cultured for 12 h.

### Plasmid construction

Plasmid construction was performed as reported previously (Tan et al., 2019). For the overexpression assay and the localization assay, the expression vector and fluorescence-labeled vector were constructed. The targeting sequences were amplified by the cDNA library of Huh7 cells using KOD-Plus-Neo DNA polymerase (#KOD-401, TOYOBO, Shanghai, China). After gel extraction, the fragment was cloned into the digested pcDNA3.1 vector (digestion sites, HindIII and BamHI) using a seamless cloning kit (#C112-01, ClonExpress II One Step Cloning Kit, Vazyme, Nanjing, China).

### qPCR assay

Real-time PCR was performed as reported previously (Tan et al., 2019) using the QuantStudio 6 Flex Real-Time PCR System (ABI, Thermo Fisher, Shanghai, China) and Roche LightCycler® 480 (Roche, Switzerland), and the following PCR program: denaturation at 95 °C for 10 min; amplification for 45 cycles at 95 °C for 15 s; annealing and extension at 60 °C for 1 min. Two × SYBR Green qPCR Master Mix (#B21203, Bimake, Shanghai, China) were used for the detection of RT-qPCR. Specific amplifications for certain PCR reactions were assessed using a melting curve. One negative control reaction, in which the cDNA template was replaced by water, was performed to avoid potential contamination. The sample from each well was repeated three times, and the comparative Ct (2−ΔΔCt) value method was used for relative quantification. GAPDH (NM_002046.6) was used as the reference gene.

### Silver stain

Silver stain was performed according to the manual (# P0017S, Beyotime, Nanjing, China). Briefly, the page glue was first fixed with ethanol/acetic acid fixing solution at room temperature for 1 h. After that, it was washed with 30% ethanol and deionized water. After sensitization and water washing, silver staining was performed. After 1.5 minutes of washing with water, color developed, and the reaction was terminated after the expected band was observed.

### Immunoprecipitation and co-immunoprecipitation

Immunoprecipitation and nuclear co-precipitation were performed according to the following instructions (Protein A/G beads, # B23201, Bimake, Shanghai, China). An antigen sample is first prepared. The culture medium is removed and washed with 1 × PBS twice at a ratio of 150 μL per 1.0 × 105 cells; the cells are scraped with a cell scraper and collected in a 1.5 mL EP tube. For every 20–30 μL per 1.0 × 105 cells, add binding buffer and protease inhibitor at the same ratio, mix, and place on ice for 10 min; collect the supernatant by centrifugation (4 °C, 14,000× g, 10 min), and keep on ice for later use. Vortex the magnetic beads for 1 minute to fully suspend them; take 25–50 μL of the magnetic bead suspension into a 1.5 mL EP tube. Add 200 μL binding buffer for washing and magnetic separation (place the centrifuge tube on a magnetic stand and let it stand for 2 minutes or wait for the magnetic beads to adhere to the tube wall). Aspirate and discard the supernatant. Repeat the washing once, insert the magnetic strip, magnetically separate and aspirate the supernatant. Add 200 μL of binding buffer to resuspend the beads for later use. Prepare antibody working solution, dilute the antibody sample with binding buffer, prepare antibody working solution with a final concentration of 5–50 μg/mL, and keep on ice for later use. Magnetically separate the pre-treated magnetic bead suspension and aspirate and discard the supernatant; add 200 μL of antibody working solution, resuspend quickly and place it in a mixing mixer at room temperature or gently invert the EP tube manually, and perform magnetic separation after 15 min. Collect the supernatant and place on ice for subsequent testing. Remove the magnetic strip, add 200 μL of binding buffer to the EP tube, and gently pipette to disperse the magnetic bead–antibody complex evenly. Insert the magnetic strip and perform magnetic separation. Aspirate and discard the supernatant. Repeat washing once. Add 200 μL of the antigen sample prepared in step 1, and gently pipette to disperse the antigen and magnetic bead–antibody complex. Place in an inversion mixer or gently invert the EP tube by hand and react at 4 °C overnight. Magnetically separate the magnetic bead–antibody–antigen complexes for antigen adsorption described above, collect the supernatant and place on ice for subsequent detection. Remove the magnetic strip, add 200 μL of washing buffer to the EP tube, and gently pipette to disperse the magnetic bead–antibody–antigen complex uniformly. Insert the magnetic strip and magnetically separate it. Aspirate and discard the supernatant. Repeat the washing two times. Remove the magnetic strip, add 200 μL of washing buffer, transfer the magnetic bead–antibody–antigen complex suspension to a new 1.5 mL EP tube, insert the magnetic strip and perform magnetic separation, and aspirate the supernatant. Finally, after the antigen is eluted, and the magnetic strip is drawn out, 20–50 μL of 1 × SDS-PAGE loading buffer is added to the mixture, and the mixture is heated at 95 °C for 5 minutes. Insert the magnetic strip and perform magnetic separation to collect the supernatant for SDS-PAGE detection.

### Protein model analysis

The structure of DLD was visualized using Pymol (www.schrodinger.com/pymol). The PDB accession number is 6I4Q (Szabo et al., 2019). The visualization methods were according to the instruction manual for Pymol.

### Measurement of phospholipids

This project used a non-targeted lipid histology platform based on the UPLC-MS/MS (QE Plus™) liquid chromatography-mass spectrometry system. The platform is combined with LipidSearch™ software (Thermo Scientific™) for lipid identification and pre-processing data. For each sample, 1 mL of MTBE (Methyl tert-butyl ether) was added with 400 μL of aqueous methanol (3:1, v/v), mixed well, and shaken for 60 min at 4°C. 1, v/v), mix well, and shake for 60 min at 4°C; then sonicate for 15 min in an ice bath, and add 250 μL water, mix well, and let stand at room temperature for 10 min; centrifuge at 20,000g for 5 min at 4°C. Using a glass syringe the top layer (lipid layer) was added to a clean tube using a glass syringe. The precipitated proteins were lysed with 200 μL of SDT lysate BCA quantification was then performed and the relative amounts of lipids in different samples were extrapolated based on the amount of protein in each sample. The relative amounts of lipids in different samples were then extrapolated based on the amount of protein in each sample, and then equal amounts of lipids were taken for each sample and blown dry using nitrogen. The lipid samples were dried and stored the samples were stored in a refrigerator at −80°C. For each sample, 100 μL of dichloromethane/methanol solution (1:1, v: v) was added for the mass spectrometry analysis. v: v), centrifuged, and the supernatant was taken into the sample for analysis. The samples were separated on an UHPLC system Ultimate 3000 (Thermo Scientific). The column temperature was 45 °C; the flow rate was 300 μL/min; the injection volume was 5 μL. The detection was performed in positive and negative ion mode by electrospray ionization (ESI), respectively. The samples were separated by UHPLC The samples were analyzed by mass spectrometry using a Q Exactive plus mass spectrometer (Thermo Scientific™). The software LipidSearch 4.1.30 (Thermo Scientific) was used for lipid identification (secondary identification) and quantification and other processing.

### Statistical analysis

All quantitative experiments were evaluated for statistical significance using the software GraphPad Prism v.5.0 (GraphPad Software, Inc., 7825 Fay Avenue, Suite 230, La Jolla, CA, 92037, USA). Because the sample size in some experiments was small, Wilcoxon–Mann–Whitney nonparametric tests were employed. The statistical significance (p-value) is indicated (* pL<L0.05; ** pL<L0.01).

### Illustration tool

The graphical images were created with the BioRender software (https://biorender.com/).

## Results

### 1. The nLDs exit the nucleus instead of being degraded

Past research has demonstrated that overconsumption of unsaturated fatty acids induces formation of nuclear lipid droplets (nLDs) (Romanauska and Koehler, 2021). To investigate this, we exposed Huh7 cells to 400 µM oleic acid and employed BODIPY staining to label LDs, thereby uncovering an augmented presence of nuclear LDs (Figure 1A). We then halted the oleic acid exposure and substituted it with fresh medium, allowing us to explore the metabolic fate of nuclear LDs via live cell imaging. Intriguingly, we found that the nuclear LDs appeared to exit the nucleus directly rather than undergoing gradual degradation (Figure 1B, Supplementary Video S1).

**Figure 1.**
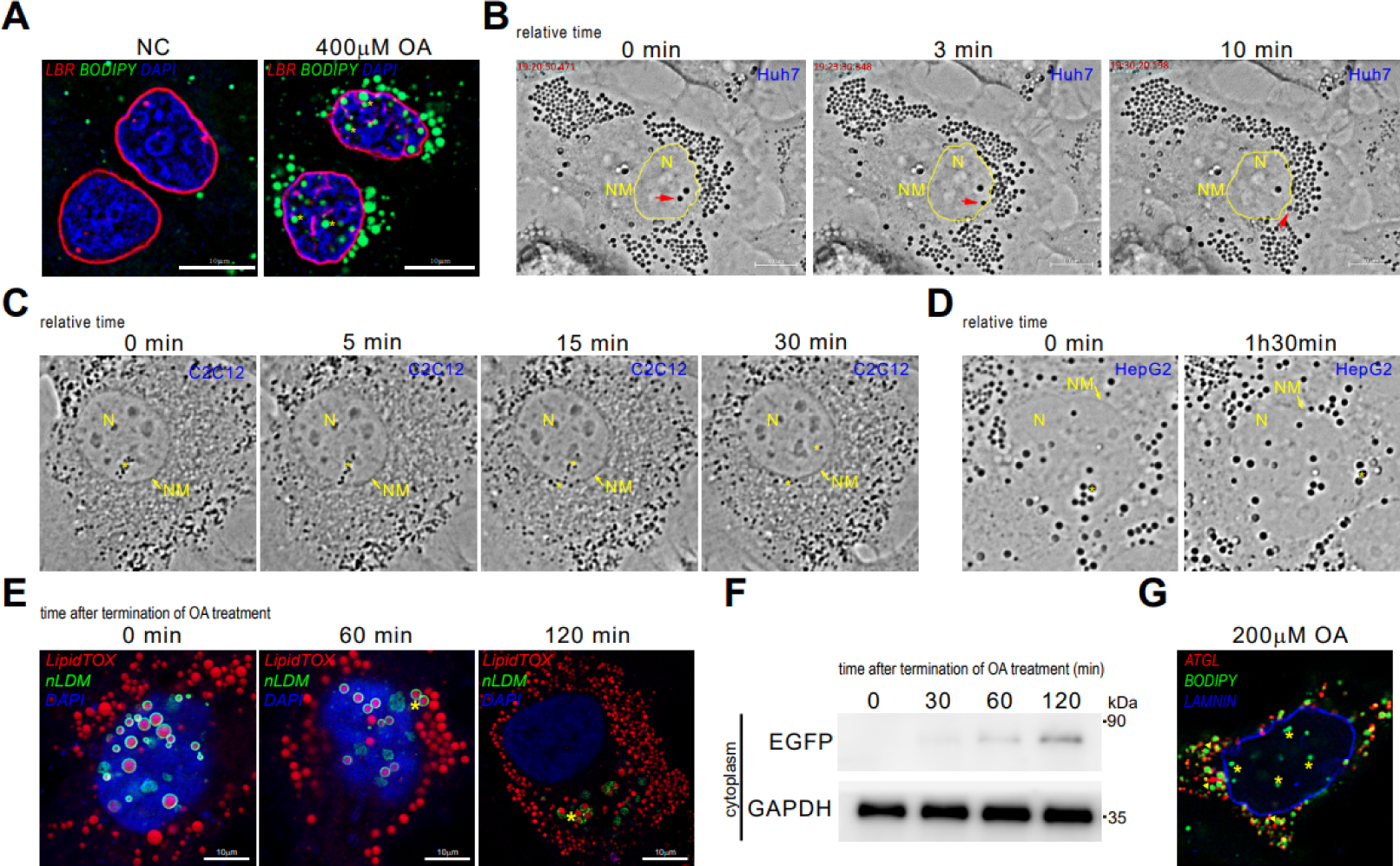
The nLDs exit the nucleus instead of being degraded. A. 400mM oleic acid treatment induced nLDs formation. red, LBR, inner nuclear membrane; green, BODIPY, lipid droplet; blue, DAPI, nucleus. B-D. real-time observation of lipid droplets in Huh7, C2C12, HepG2, respectively. N: nucleus, NM: nuclear membrane. The arrow marks the nLDs going out nucleus. E. labeling nLDs with specific vector, nLDM. F. detection of EGFP level in cytoplasm at 0, 30, 60 and 120min after oleic acid treatment. Asterisk marks the nLDs out of nucleus. G. detection of ATGL (red) localization. Asterisk marks nLDs.

To validate this observation, we utilized the EGFP fluorescent vector to specifically label nLDs, capturing their presence in the nucleus through laser confocal microscopy (Figure 1C, D, Supplementary Videos S2, S3). The detection of LDs with EGFP fluorescence signal in the cytoplasm substantiated their nuclear origin (Figure 1E). Further support was provided by western blotting, which revealed EGFP protein expression in the cytoplasm, confirming the ability of nuclear LDs to exit the nucleus (Figure 1F).

In an effort to elucidate the need for nuclear LDs to be excreted, we employed immunofluorescence to label the lipase ATGL, only to find that this lipase is not distributed on the surface of the nuclear LDs (Figure 1G). This indicates that nuclear LDs cannot undergo glycerol ester hydrolysis, unlike their cytosolic counterparts. Additionally, given the absence of mitochondria and lysosomes within the nucleus, none of the traditional LD breakdown pathways are plausible.

### 2. The accumulation of nLDs is associated with phosphatidylcholine synthesis pathway

To further dissect the mechanisms driving nLD accumulation, we conducted a series of experiments. Initially, cells were treated with choline-free medium, 400 µM oleic acid medium, and choline-free medium containing 400 µM oleic acid, after which the number of nLDs was quantified. Notably, we observed a significant surge in nLDs in the presence of both choline-free medium and 400 µM oleic acid (Figure 2A, B), establishing this as a methodology for inducing nLD generation.

**Figure 2.**
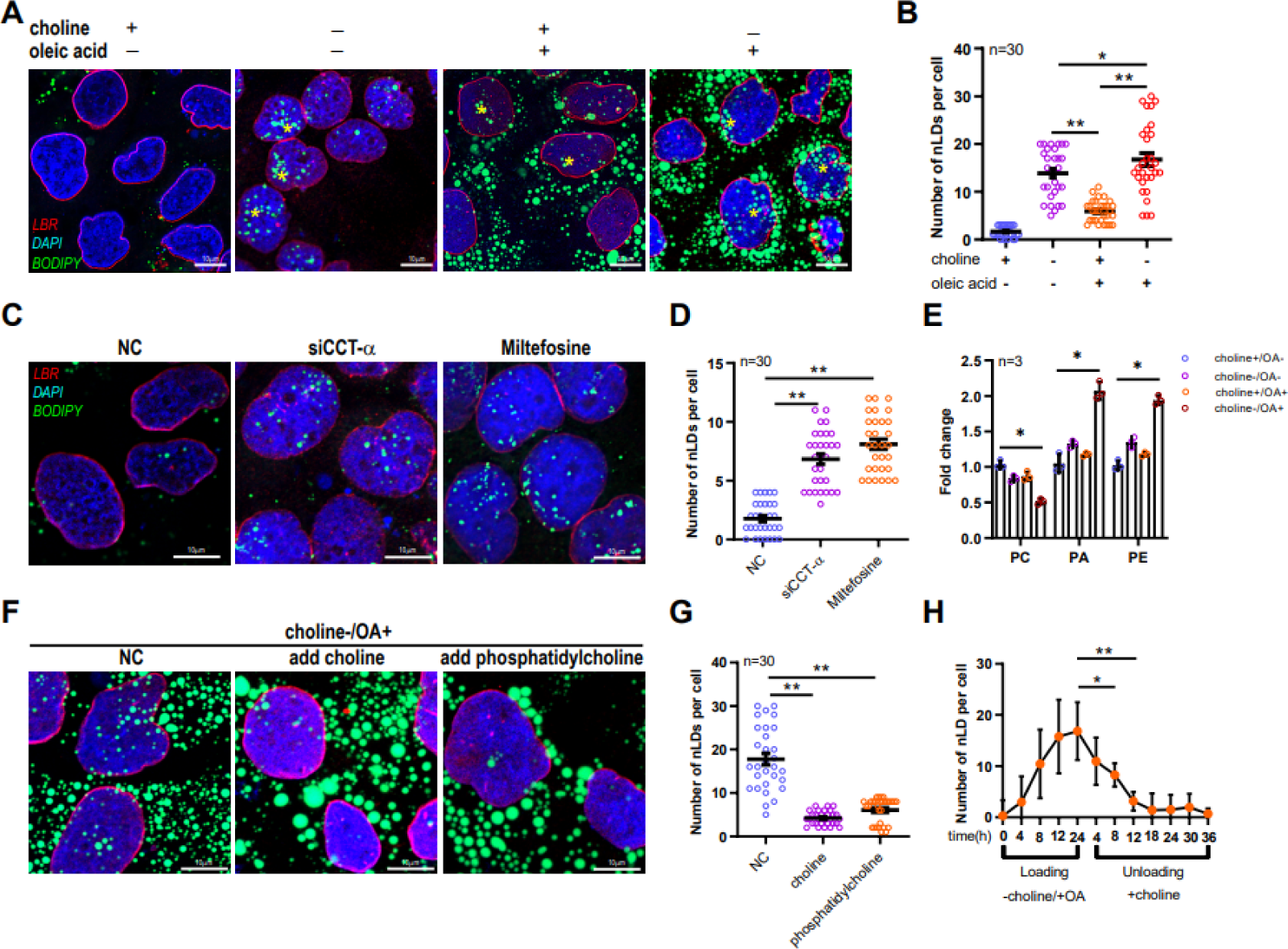
The accumulation of nLDs is associated with phosphatidylcholine synthesis pathway. A-B. Cells were treated with oleic acid, choline-restricted and oleic acid combined with choline-restricted, respectively, to detect the number of nLDs. C-D. Cells were interfered with CCT-a and treated with miltefosine to detect the number of nLDs. E. Cells were induced to produce nLDs and the content of each phospholipid fraction was detected. F-G. Cells were treated with choline-restricted medium to restore choline supply and the number of nLDs was detected. H. nLDs number was analyzed without bias by High content screening laser confocal microscopy. *, p<0.05, **, p<0.01.

Additionally, we executed RNA interference of CCT-α and utilized miltefosine to inhibit the phosphatidylcholine synthesis pathway. These interventions resulted in a pronounced increase in the quantity of nLDs (Figure 2C, D), alongside a corresponding reduction in phosphatidylcholine levels and elevation in PA and PE levels (Figure 2E).

Probing the role of phosphatidylcholine in nLD accumulation, we evaluated the number of nuclear LDs post-restoration of choline supply or after external PC addition, following treatment with choline-free medium and 400 µM oleic acid. Interestingly, upon restoring the choline supply, we noticed a decrease in nuclear LDs (Figure 2F, G), thereby underscoring the role of phosphatidylcholine in nLD generation. This finding was further solidified by high-intensity unbiased confocal microscopy, which depicted a statistically significant reduction in nLD count upon choline replenishment (Figure 2H).

### 3. The nLDs out of nucleus are activated by RhoA signaling

To delve deeper into the signals prompting nLD egress from the nucleus, we analyzed mRNA expression profiles at four distinct time points before, during, and after the induction of nLD formation. Gene Ontology (GO) and pathway analyses of the transcriptomic data revealed substantial variations in cell cycle-associated signals. Choline-deficient treatment led to the downregulation of cell cycle-associated genes and suppression of relevant signaling pathways (Table 1 and Figure 3A).

**Figure 3.**
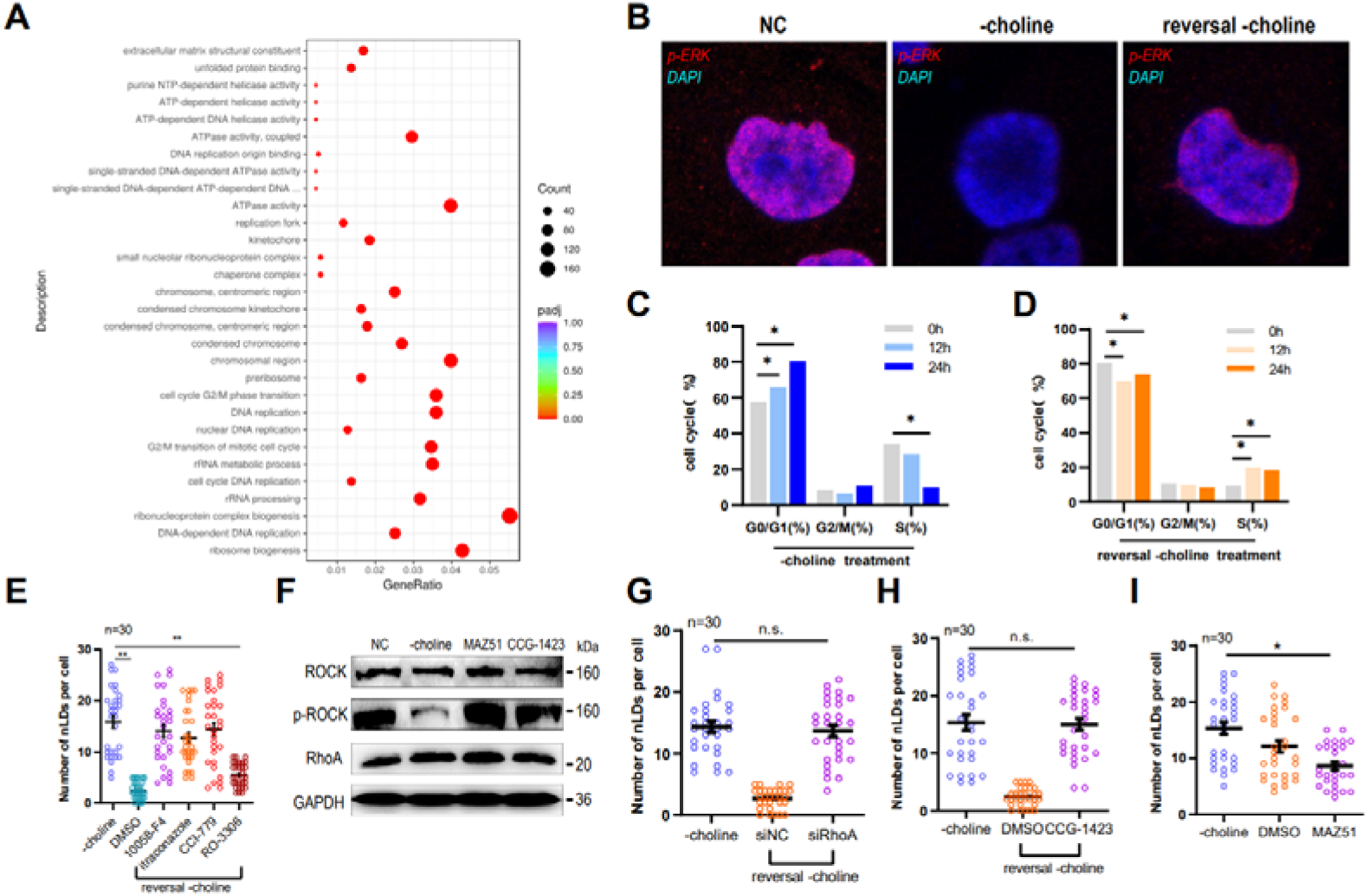
The nLDs out of nucleus are activated by RhoA signaling. A. High-throughput RNA sequencing to detect the signaling pathways that change before and after the induction of nLDs generation. B. Detection of ERK phosphorylation levels during the induction of nLDs generation and during the induction of nLDs exit from the nucleus. C. Flow cytometry to detect the number of cells at each stage of mitosis during the induction of nLDs generation. D. Flow cytometry to detect the number of cells at each stage of mitosis during the induction of nLDs exit from the nucleus. E. Induction of nLDs to induce nLDs exit, cells were arrested in G0, G1, and G2 phases using 10058-F4, itraconazole, CCI-779, and RO-3306 cell cycle inhibitors, respectively, and the number of nLDs was detected. F. Cells were treated with RhoA activator MAZ51 and inhibitor CCG-1423 to detect the level of downstream signal ROCK phosphorylation, respectively. G. Induction of nLDs out of the nucleus, cells were treated with RhoA interference fragment and the number of nLDs was detected. H-I. Induction of nLDs out of the nucleus, cells were treated with CCG-1423 and MAZ51, respectively, and the number of nLDs was detected. N=30, *, p<0.05, **, p<0.01.

**Table 1.**
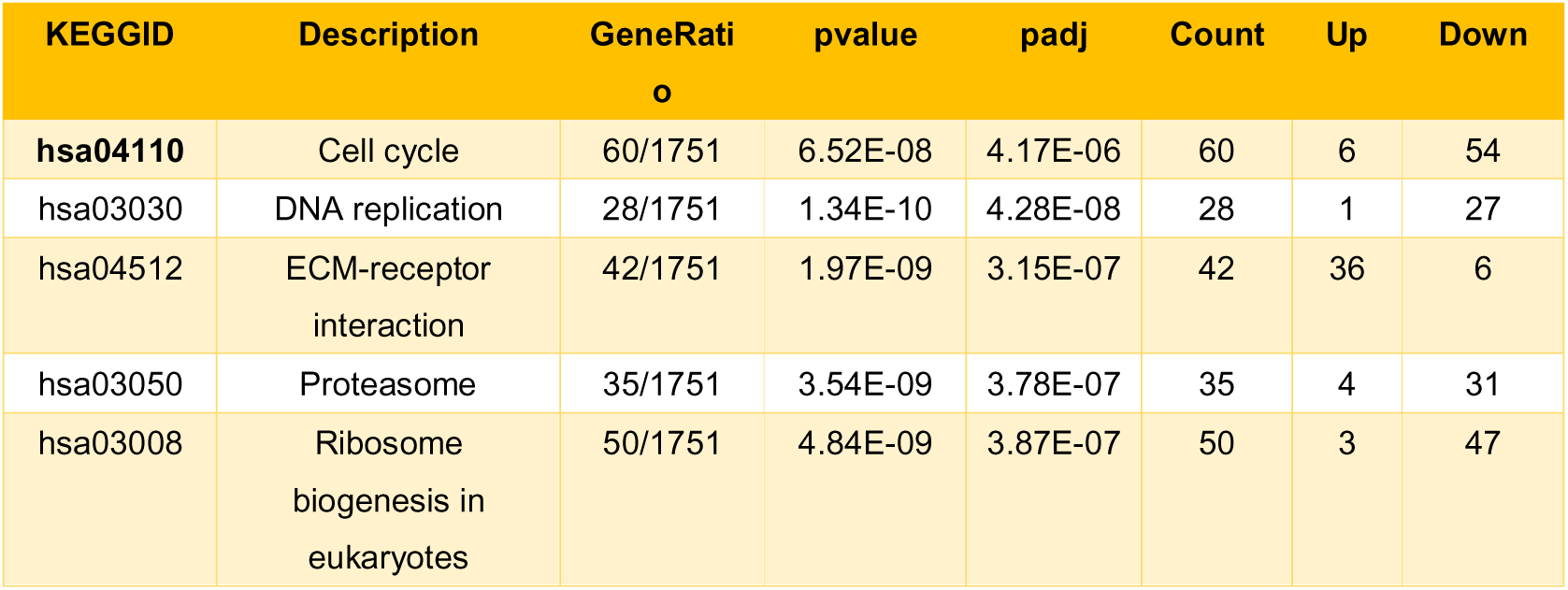
Pathway Inhibition with Choline-Restricted Treatment. The table shows the pathways inhibited by choline-restricted treatment. The KEGG IDs, pathway descriptions, gene ratios, p-values, adjusted p-values (padj), and the count of upregulated (Up) and downregulated (Down) genes are provided.

Immunofluorescence measurements of phosphorylation levels of cell cycle regulators, such as ERK, indicated that choline deprivation results in decreased p-ERK levels, while resupplying choline elevates p-ERK levels (Figure 3B). Flow cytometry analyses demonstrated that choline deprivation leads to an increased population of cells in the G0/G1 phase and a decreased S phase fraction (Figure 3C). Conversely, following choline resupply, the number of G0/G1 phase cells diminished while S phase cells increased (Figure 3D).

To assess the influence of cell cycle signals on nLD egress, we arrested cells in the G0/G1, S, or G2 phase using various inhibitors post nLD induction, and quantified the nLDs. The data demonstrated a significant reduction in nuclear LDs during the G2 phase (Figure 3E), implying that the G2 phase commencement has a vital impact on nLD nucleation.

Building on literature and our transcriptomic data, we hypothesized that choline deprivation might affect RhoA signaling, thereby influencing downstream protein phosphorylation. This was corroborated by a Western blotting assay, which revealed that choline deprivation suppressed RhoA signaling (with RhoA activator MAZ51 and inhibitor CCG-1423 as controls) (Figure 3F).

To examine the role of RhoA signaling on nLD nucleation, we used RNA interference (RNAi) to suppress RhoA mRNA expression post nLD induction, and subsequently quantified the nLDs. The results revealed that RhoA expression inhibition led to suppressed nLD nucleation (Figure 3G). Additionally, following nLD induction, we treated cells with the RhoA inhibitor CCG-1423 and quantified nLDs post nLD egress induction. The results indicated that RhoA activity inhibition led to suppressed nLD egress (Figure 3H). Moreover, upon introducing a RhoA activator to choline-deprived cells and assessing nLD count, we found a significant reduction in nLDs post RhoA activator addition compared with the control group (Figure 3I). Collectively, these findings imply that RhoA signaling regulates the egress of LDs from the nucleus.

### 4. The movement of nLDs is activated by Rab8a derived from INE

To further elucidate the process of nLD egress from the nucleus, we utilized a live-cell workstation to dynamically observe nLDs post their formation and subsequent nuclear exit. Prior to inducing nLD exit, nLDs primarily remained stationary, with occasional minor vibrations in situ (Supplementary Video S4), whereas post nLD exit induction, nLD motion intensified (Supplementary Video S5).

To explore the role of Rab8a in this process, we assayed its expression levels in the nucleus and cytoplasm before and after LD exit induction using Western blotting. Overall, Rab8a expression remained relatively constant across the two conditions (Figure 4B). However, following nuclear isolation and nLD extraction, measurements of Rab8a levels on the nLD surface demonstrated a significantly heightened expression of Rab8a post nLD exit induction (Figure 4C, D).

**Figure 4.**
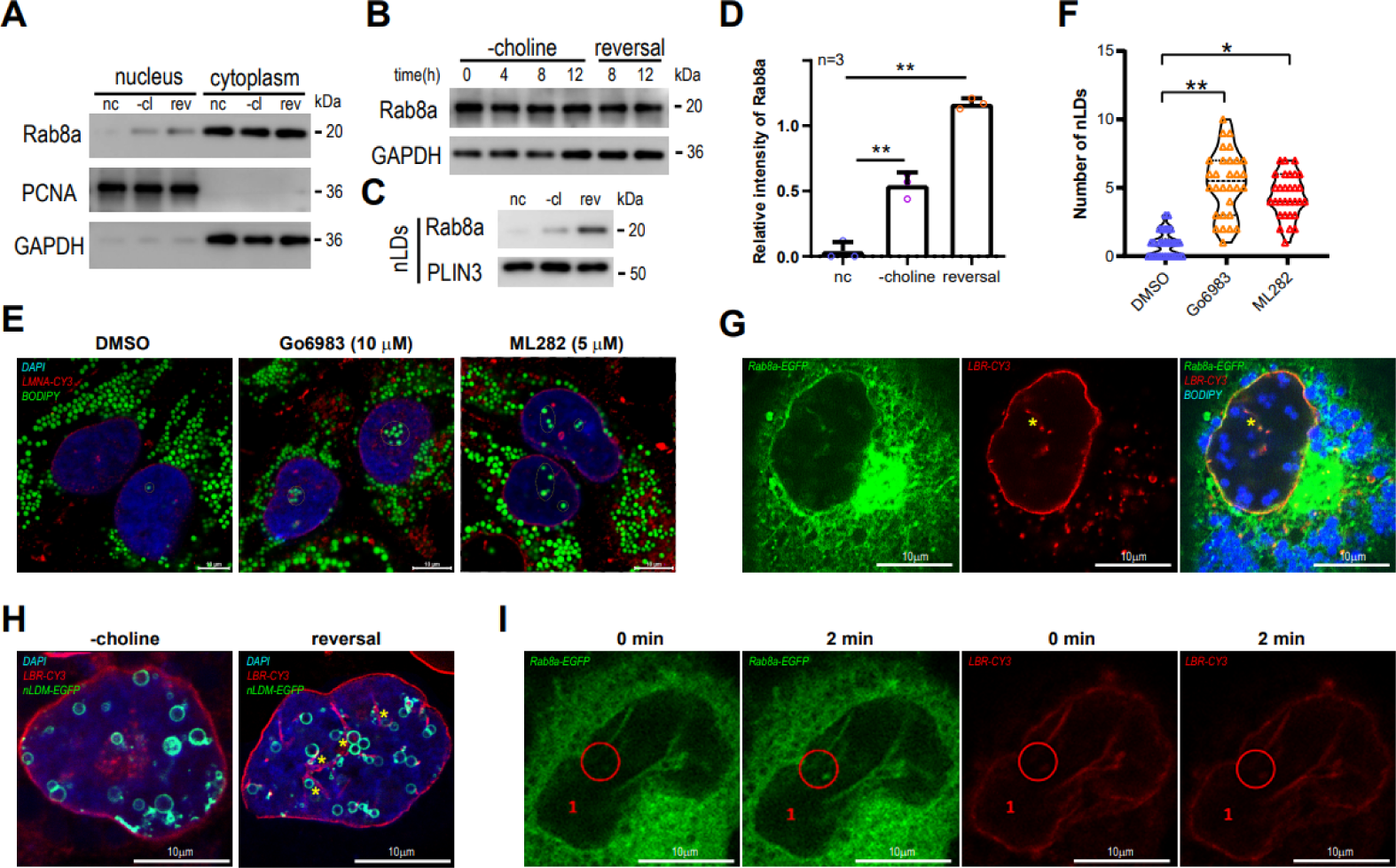
The movement of nLDs is activated by Rab8a derived from INE. A. Rab8a levels were detected in the nucleus and cytoplasm during the induction of nLDs production and efflux phases, respectively. B. Total Rab8a levels were detected in the cells during the induction of nLDs production and efflux phases. C-D. Lipid droplets in the nucleus were isolated during the induction of nLDs production and efflux phases, and their surface Rab8a levels were detected. E-F. Cells were treated with RhoA inhibitor and activator, respectively, to induce nLDs efflux, and the number of nLDs was detected. The number of nLDs was detected. G. Immunofluorescence detection of Rab8a localization on the type INE. H. Induction of nLDs exiting the nucleus was accompanied by an increase in the number of INE and contact with nLDs. I. Fluorescence bleaching recovery assay to detect the mobility of Rab8a in the INE.

To further scrutinize Rab8a’s impact on nLD motility, we overexpressed NLS-Rab8a (Rab8a with a nuclear localization signal) and employed a live-cell workstation to monitor nLD motility. Our observations indicated that Rab8a overexpression stimulated nLD motility (Supplementary Video S6). Moreover, treating cells with a Rab protein inhibitor to induce nLD nuclear egress led to a significant increase in nuclear LDs in the inhibitor-treated group (Figure 4E, F), signifying Rab protein’s regulatory role in nuclear LD movement.

Investigating the origin of Rab8a on the nLD surface, we labeled Rab8a with Rab8a-EGFP and used laser confocal detection. Our observations indicated abundant Rab8a expression in the nuclear membrane and the inner nuclear reticulum (INR) (Figure 4G). This led us to postulate that Rab8a on the nLD surface within the nucleus stemmed from the INR. Immunofluorescence labeling of LBR protein-reactive INR locations and subsequent co-localization analysis with nLDs supported our hypothesis, revealing increased INR and nLD contact post nLD nuclear exit induction (Figure 4H). Additionally, we employed a fluorescence recovery after photobleaching assay to confirm the mobility of Rab8a in the INR, which demonstrated a gradual recovery of the fluorescence signal on the INR over time (Figure 4I and Supplementary Video S6).

### 5. The nLDs exit nucleus through the gap of lamina induced by LMNA S392 phosphorylation

The nuclear envelope, comprised of the outer nuclear membrane, inner nuclear membrane, and lamina, consisting of dense, neutral fibrils, maintains the cytoskeleton. We initiated nLD nuclear egress by immunofluorescence labeling of the lamina with LMNA and extranuclear membrane with NUP98, followed by laser confocal microscopy detection. We observed an incomplete lamina with noticeable gaps at the site of nLD and nuclear membrane contact, while the extranuclear membrane remained intact (Figure 5A). LMNA depolymerization was associated with its phosphorylation level.

**Figure 5.**
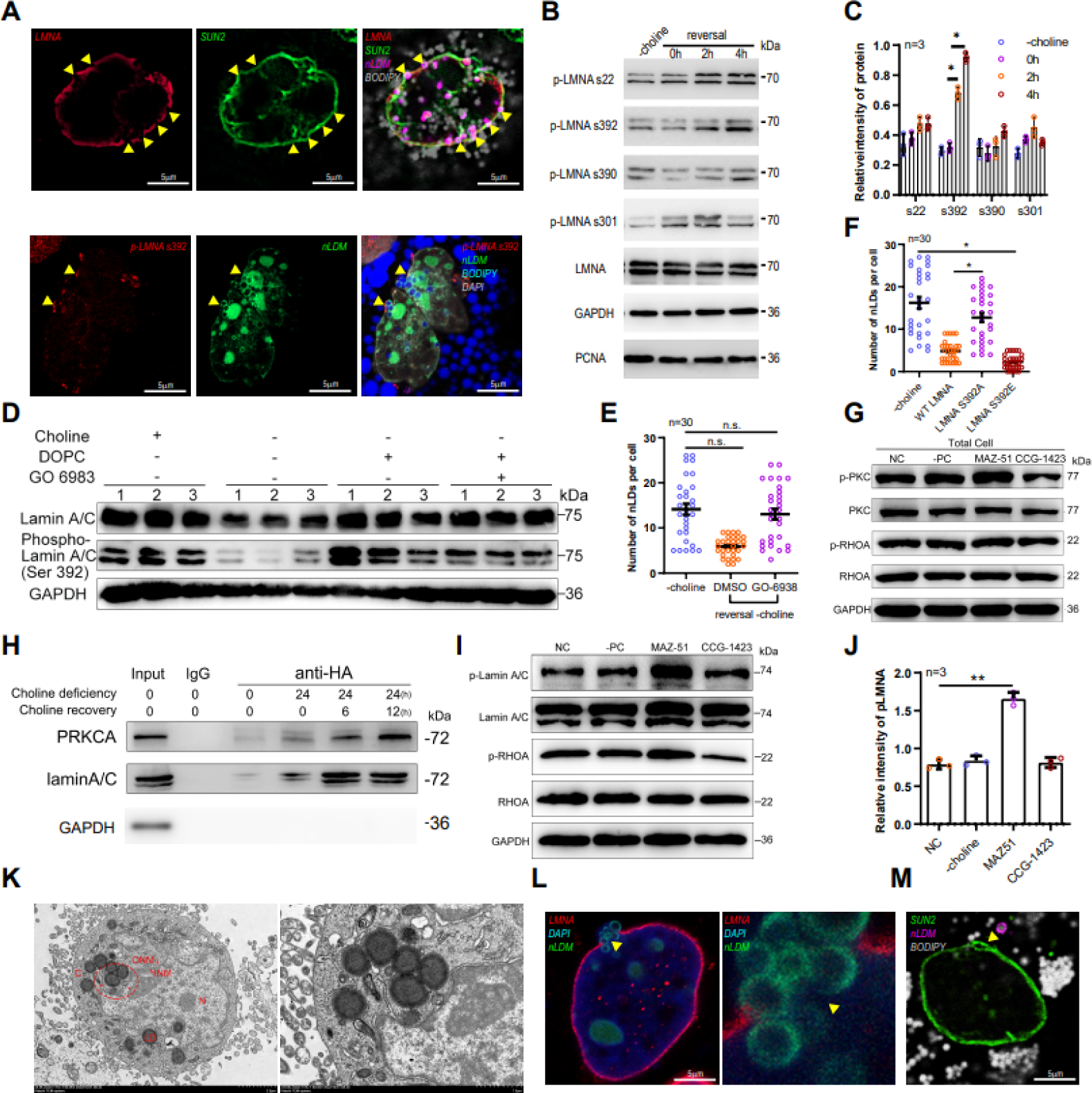
The nLDs exit nucleus through the gap of lamina induced by LMNA S392 phosphorylation. A. Immunofluorescence labeling of lamin, nuclear outer membrane (SUN2) and phosphorylated lamin S392 to detect protein distribution at the nLDs contact site. B-C. Induction of nLDs exit from the nucleus to detect LMNA site-specific phosphorylation levels. D. Detection of LMNA s392 phosphorylation levels using the cell cycle inhibitor GO6983. E. Treatment of cells with GO6983 to induce nLDs exit from the nucleus to detect nLD numbers. F. Construction of s392 inactivation and sustained activation mutants to detect nLD numbers. G. Detection of PKC phosphorylation levels using RhoA inhibitor and activator-treated cells. H. Detection of PKC binding capacity to LMNA upon induction of nLDs out of the nucleus using the Co-IP technique. I-J. Detection of LMNA phosphorylation levels using RhoA inhibitor and activator-treated cells. K. Transmission electron microscopy to detect nuclear membrane morphology near the nuclear sites. L-M. Immunofluorescence to detect nuclear membrane morphology near the nuclear sites. *, p<0.05, **, p<0.01.

By using Western blotting to assess phosphorylation levels at LMNA S22, S390, and S392 sites, we found that only S392 showed an elevated expression level following nLD nuclear exit induction (Figure 5B, C). Laser confocal observation of LMNA S392 immunofluorescence labeling revealed a strong fluorescent signal at the contact site between nLD and the nuclear membrane (Figure 5D).

Previous research suggests a pivotal role of PKC in regulating LMNA phosphorylation. When cells were treated with the PKC inhibitor GO-6983, a significant reduction in S392 phosphorylation was observed (Figure 5E). Assaying nLD nuclear egress in cells treated with the PKC inhibitor showed inhibition (Figure 5F). Following LMNA expression interference using RNAi, we constructed and expressed LMNA S392A and S392E mutants and assessed nLD egress. LMNA S392A expression inhibited nLD egress, while LMNA S392E expression promoted it (Figure 5F).

We investigated the effect of RhoA signaling on PKC and LMNA S392 phosphorylation by treating cells with a RhoA inhibitor, and assessing PKC expression and phosphorylation levels. PKC phosphorylation significantly reduced, while its expression remained stable (Figure 5G). Using immunoprecipitation, we found that inhibiting RhoA signaling diminished the binding of LMNA to PKC (Figure 5H). Inhibition of RhoA also considerably decreased LMNA S392 phosphorylation (Figure 5I, J).

We utilized transmission electron microscopy to observe the detailed structure of the LD exit site, revealing a discontinuous nuclear membrane near the LD exit site, with an inner nuclear membrane separating the nucleus (Figure 5K). This structure was also discerned at the LD exit site by immunofluorescence labeling (Figure 5L, M), consistent with our electron microscopy results.

### 6. The nLDs exit nucleus carrying with DNA fragments

The nucleus, acting as the storage for an organism’s genetic material, contains DNA. Earlier studies showed that histone binding receptors on the surface of bacterial lipid droplets (LDs) can anchor DNA, offering protection to the genetic material. We hypothesized that nuclear lipid droplets (nLDs) in mammalian cells might transport DNA out of the nucleus. To validate this, we extracted nLD samples and performed surface DNA content analysis, which revealed DNA fragments of different sizes on the LD surface (Figure 6A).

**Figure 6.**
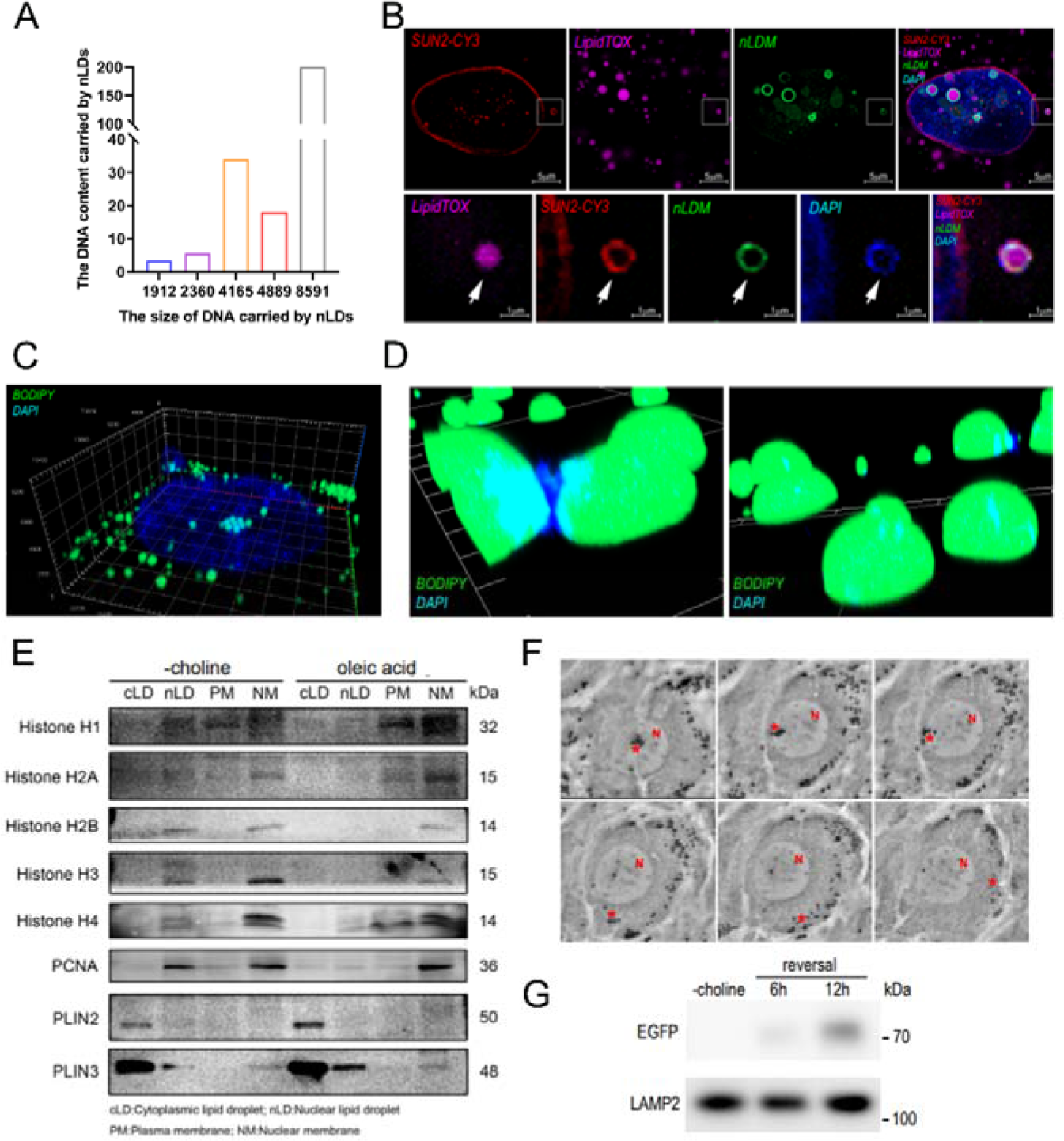
The nLDs exit nucleus carrying with DNA fragments and are degraded by autophagy. A. The DNA content carried by nLDs. B. Immunofluorescence labeling of nLDs was found to carry DNA after nucleation. C. Laser confocal 3D scanning was performed to detect the distribution of nLDs and DNA. D. Purified nLDs were stained using DAPI and found to carry DNA. E. WB detection of nLDs carrying histones. F. Live cell observation was performed to detect the dynamics of nLDs after nucleation.G. Lysosomes were extracted and WB detection of EGFP content.

We further labelled nLDs and DNA with DAPI. The findings showed DAPI fluorescence signals on the surface of LDs that had exited the nucleus, indicating the existence of DNA on these LD surfaces (Figure 6B). Laser confocal 3D scanning revealed contact between nLDs and DNA (Figure 6C). After extracting the nuclei to isolate the nLDs and incubating them with cellular DNA in vitro, we thrice eluted them, labelled the LDs with DNA, and found co-localization of LDs and DNA through laser confocal detection (Figure 6D).

To confirm the ability of nLDs to bind DNA, we extracted the nuclei to isolate the nLDs and detected the expression of histones H1, H3, and H4. The findings showed substantial histone expression on the nLDs (Figure 6E), indicating their capability to bind DNA.

### 7. The nLDs are degraded by autophagy in cytoplasm

The ultimate fate of nLDs after their departure from the nucleus was observed using a live cell workstation (Figure 6F). It was found that these LDs were eventually disassembled and vanished within the cytoplasm. The degradation of cytoplasmic LDs often follows an established pathway involving autophagic lysosomes.

To further verify the binding of nLDs to autophagosomes, we tagged LDs with a specific EGFP fluorescent vector. After isolating autophagosomes, a Western blot assay was performed to detect the expression of EGFP protein. The results displayed a relatively low level of EGFP expression in the extracted autophagosomes (Figure 6G). This suggests that the breakdown of nLDs exiting the nucleus involves the autophagic process.

## Discussion

Our study unveils a novel, evolutionarily conserved mechanism for the exit of large molecular entities, including nLDs, from the nucleus. This discovery is unprecedented and of significant importance, as it challenges our understanding of cellular processes and the dynamics of the nucleus.

While the selective transportation of substances between the nucleus and cytoplasm primarily relies on nuclear pores, the size of nLDs prevents them from passing through these pores. The nuclear pore complex (NPC) is a massive protein assembly that serves as the primary gateway between the nucleus and the cytoplasm. Structurally, the NPC is composed of multiple protein subunits, collectively termed nucleoporins, which assemble to form a channel that spans the nuclear envelope (Suntharalingam and Wente, 2003). Functionally, the NPC facilitates the selective transport of RNA, proteins, and ribonucleoprotein particles, which is crucial for numerous cellular processes, including gene expression, DNA repair, and cell cycle progression (Lin and Hoelz, 2019).

However, entities like nLDs, due to their large size, are unable to traverse the NPC for transport. This implies that nLDs might possess a unique mechanism for exiting the nucleus, distinct from the conventional transport pathways mediated by the NPC(Chandra and Lusk, 2022; Tang et al., 2022). We discovered that nLDs exit the nucleus by traversing the triple layers of the nuclear membrane, involving localized depolymerization of the nuclear lamina.This mechanism shares similarities with other instances of nuclear material export, such as the exit of herpes viruses and large ribonucleoprotein (RNP) granules, suggesting a general mechanism for the export of large molecules from the nucleus (Mühlbauer et al., 2015; Myllys et al., 2016; Roller et al., 2014; Seo et al., 2013; Speese et al., 2012).

LDs, a multifunctional organelle with a unique structure, have gained attention from cell biology researchers. The presence of nLDs in various species, including yeast, nematode, mouse, and human cells (Jin et al., 2020; R and P, 2013; Romanauska and Koehler, 2021; Romanauska and Kohler, 2018; Soltysik et al., 2019; Soltysik et al., 2021), has sparked interest in their origin and functions. Notably, the number of nLDs significantly increases in specific environments, such as in hepatocytes of mice with NAFLD (Ostlund et al., 2022; Shin et al., 2019). Recent studies have shown that the intranuclear membrane synthesizes lipids and serves as the site of nLD generation (Romanauska and Kohler, 2018). The generation process of nLDs is similar to cytoplasmic LDs, involving glyceride synthesis, aggregation, lens structure formation, budding, and release from one side of the bilayer membrane (Romanauska and Kohler, 2018). Seipin protein mediates the interaction of LDs with the phospholipid membrane. It has been found that LD formation in the nucleus is associated with lipid metabolic stress in the nuclear membrane, as LDs shield it from unsaturated fatty acid stimulation (Romanauska and Kohler, 2021). Furthermore, nLDs regulate cellular phosphatidylcholine synthesis. Under tunicamycin stimulation, endoplasmic reticulum stress occurs, inhibiting phospholipid synthesis and leading to an increase in nLDs, which recruit phosphatidylcholine synthase and relieve endoplasmic reticulum stress (Soltysik et al., 2019). After fulfilling their function, the fate of nLDs is of interest. Similar in size to cytoplasmic LDs, with a diameter of approximately 500 nm to 1 μm, nLDs also occupy more space in the nucleus. Therefore, we investigated how such an organelle within the nucleus undergoes catabolism. Surprisingly, we discovered that nLDs can exit the nucleus directly instead of being disassembled and lost within the nucleus. Our findings provide insights into the ultimate destination of nLDs, completing the process of nLD formation to excretion. We observed the exit of nLDs from the nucleus in cells from various species and propose that this is a typical and normative cellular process.

The nucleus, a fundamental organelle in eukaryotic cells, houses genetic material and selectively transports molecules, such as RNA and proteins, through nuclear pores. However, the diameter of nLDs exceeds 500 nm, making it impossible for them to exit the nucleus through the nuclear pores. In our study, we observed that nLDs exit the nucleus by passing through the triple layers of the nuclear membrane, including the inner nuclear membrane, outer nuclear membrane, and nuclear lamina.This mechanism shares similarities with other instances of nuclear material export, such as the exit of herpes viruses and RNP granules, suggesting a general mechanism for the export of large molecules from the nucleus.During viral envelope expulsion, for example, protein kinase C (PKC) phosphorylates the nuclear lamina, disrupting it and allowing the viral particle bound to the inner nuclear membrane (INM) to move into the perinuclear space between the INM and outer nuclear membrane (ONM) (Lee and Chen, 2010). Subsequently, the INM surrounding the virus fuses with the ONM, releasing the naked viral shell into the cytoplasm. Similarly, RNP granules utilize a nuclear envelope budding pathway for their export, involving local depolymerization of the nuclear lamina mediated by phosphorylation of A-type lamin (Speese et al., 2012). Another example is the Frizzled Nuclear Import (FNI) signaling, which organizes nuclear fragments into ribonucleoprotein granules containing mRNAs encoding postsynaptic proteins. These DFz2C granules are localized in the perinuclear space and require the presence of A-type lamin LamC, with atypical PKC (aPKC) potentially phosphorylating LamC during the formation of DFz2C granules through invagination of the inner nuclear membrane. Disruption of this process results in a phenotype similar to laminopathy models.The mechanism of nuclear envelope exonucleation, observed in LDs, herpesviruses, and large RNP granules, involves phosphorylation of LMNA by PKC, localized depolymerization of the nuclear lamina, and subsequent expulsion into the cytoplasm. We propose that this conserved mechanism represents a general process for the exit of large molecules from the nucleus.

Our findings shed light on the potential involvement of nLDs in the metabolism of nuclear material, as evidenced by their carrying of genetic material. The presence of histones on the surface of nLDs suggests their role in DNA carriage, although the exact recruitment mechanism remains to be investigated. We propose that nLDs may play a role in clearing metabolic waste from the nucleus, including DNA fragments, thereby contributing to cellular homeostasis under stress (Zhang et al., 2017). Additionally, emerging studies have linked nLDs to cellular senescence, suggesting that their removal could have implications for lifespan extension. This connection highlights the significance of further exploring nLDs as a valuable target for research on stress and aging-related diseases.

Despite our groundbreaking observations, this study has several limitations. Firstly, our understanding of the mechanistic process for nLDs exiting the nucleus is still in its early stages, and more detailed explorations are required to fully understand the involved molecular pathways. Secondly, while our observations are consistent across multiple species, further investigation is needed to confirm if this process is universal. Lastly, the functional implications of nLDs and their potential role in metabolic waste clearance and lifespan extension are primarily speculative at this stage. These theories should be validated with experimental evidence in future studies.

Drawing from our pioneering findings, several areas of future research emerge. First, more detailed explorations of the mechanistic process for nLDs exiting the nucleus are needed. This includes understanding the precise molecular pathways involved and whether there are variations among different cell types or species. Second, the functional implications of nLDs require further investigation. In particular, their proposed roles in metabolic waste clearance and lifespan extension are significant and should be tested with experimental evidence. Third, the clinical relevance of our findings should be investigated, such as whether the process and function of nLDs contribute to the pathogenesis of diseases like NAFLD or age-related conditions. As the study of nLDs is still in its early stages, we foresee a vast field of research possibilities that can contribute significantly to our understanding of cellular processes and potential therapeutic interventions.

In summary, our investigation has unearthed a hitherto unknown, evolutionarily conserved mechanism facilitating the egress of large molecular entities, such as nLDs, from the nucleus. This seminal discovery not only challenges established paradigms in cellular processes and nuclear dynamics but also underscores the significance of our study. We have meticulously explored the biogenesis, function, and ultimate fate of nLDs, with a particular emphasis on their transit from the nucleus to the cytoplasm. The implications of our findings are profound, suggesting that nLDs play pivotal roles in cellular processes such as metabolic waste clearance and potentially lifespan extension. Our work heralds a new era of research into the role of nLDs in diverse physiological and pathological contexts, and underscores the potential of nLDs as diagnostic biomarkers and therapeutic targets. The revelation of a conserved mechanism for the export of large molecules from the nucleus represents a significant leap forward in our understanding of cellular biology. This discovery holds the potential to catalyze a paradigm shift in a wide range of biological and medical research, setting the stage for future explorations in this riveting area of study.

## Statements of approval

We confirm that all methods were performed in accordance with the relevant guidelines and regulations of Ethics Committee of Huazhong Agricultural University.

## Competing Interest Statement

The authors declare that they have no competing interests.

## Availability of Data and Materials

The data used to support the findings of this study are available from the corresponding author upon request.

## Author Contributions

Y.J. conceived and designed the experiments. Y.J., J.H., Z.C.Z. performed the experiments. Y.J., J.H. wrote the manuscript. Y.J.T performed the EM experiments. Z.Q.R. supervised this study.

## Acknowledgements

This work was supported by the National Key Research and Development Program of China (No. 2021YFF1000601), the National Natural Science Foundation of China (32172700), the Joint Funds of the National Natural Science Foundation of China (U20A2052), and the Fundamental Research Funds for the Central Universities (2662022DKPY002).

## Notes

### Competing Interest Statement

The authors have declared no competing interest.

