## Supplementary figures and images for "Nuclear Lipid Droplets Exit to the Cytoplasm: A Novel Mechanism of non-nuclear pore-dependent material exchange"

### supplementary video 1

#### Slide 1
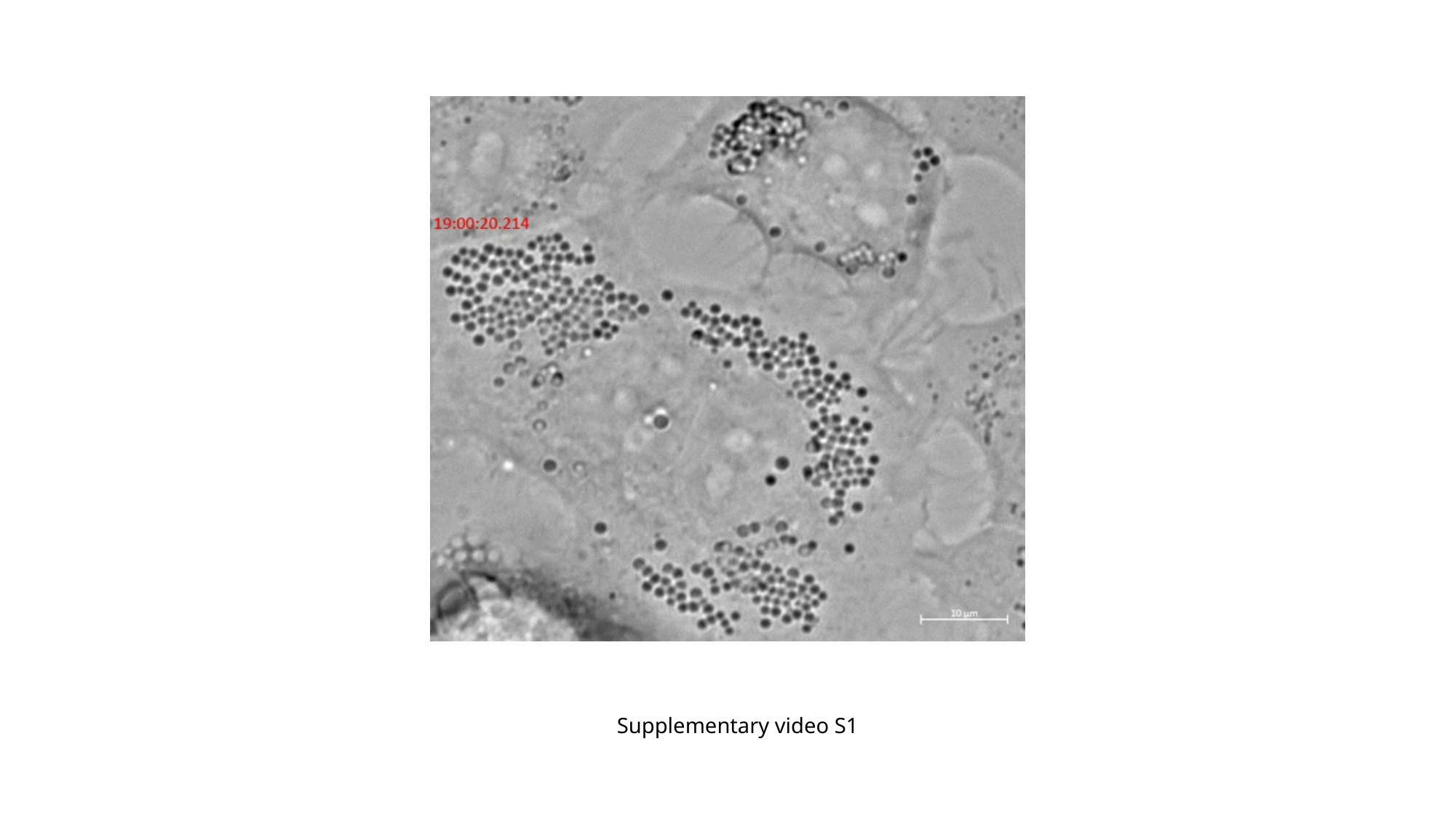

Supplementary video S1

### supplementary video 2

#### Slide 1
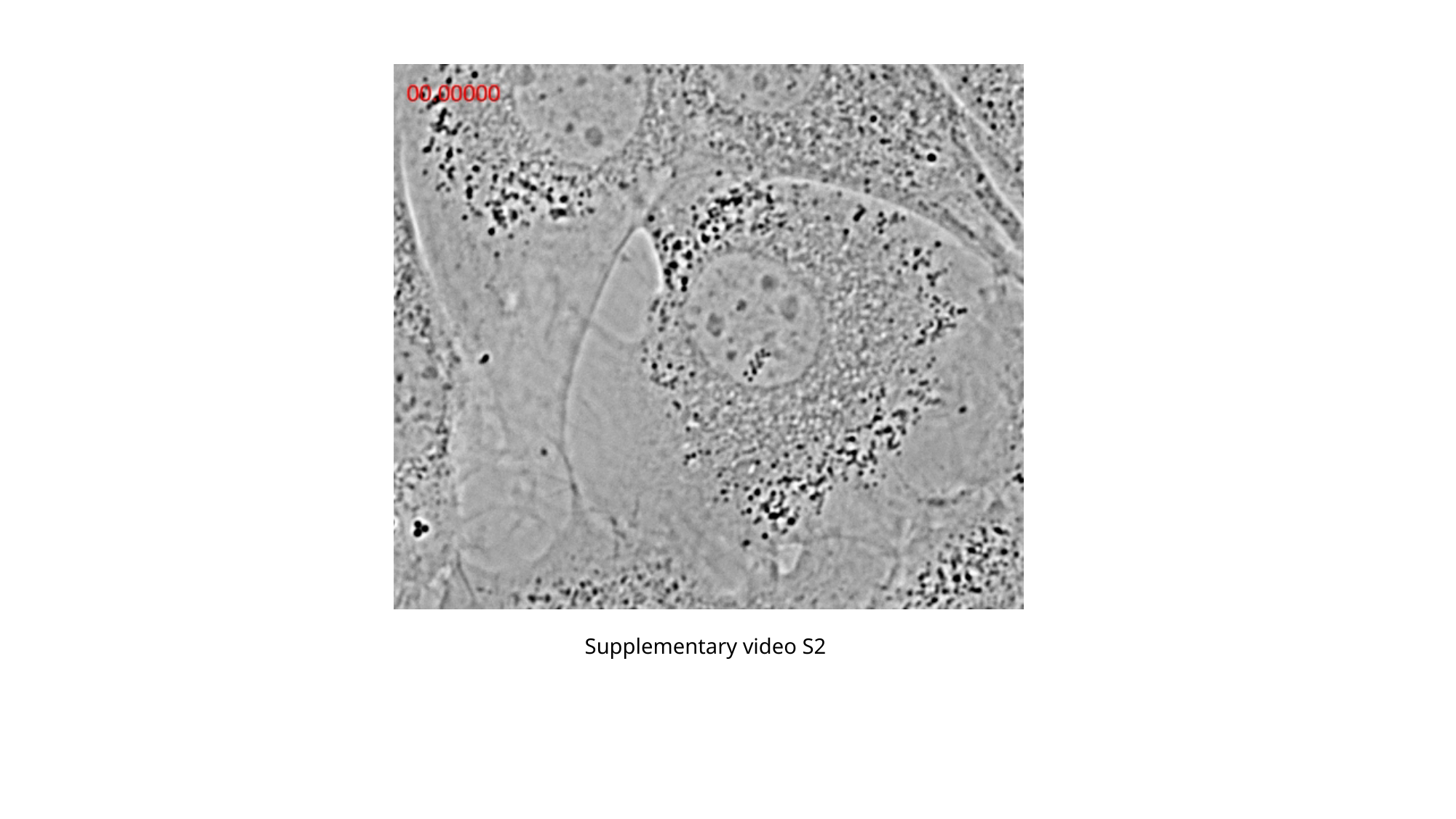

Supplementary video S2

### supplementary video 3

#### Slide 1
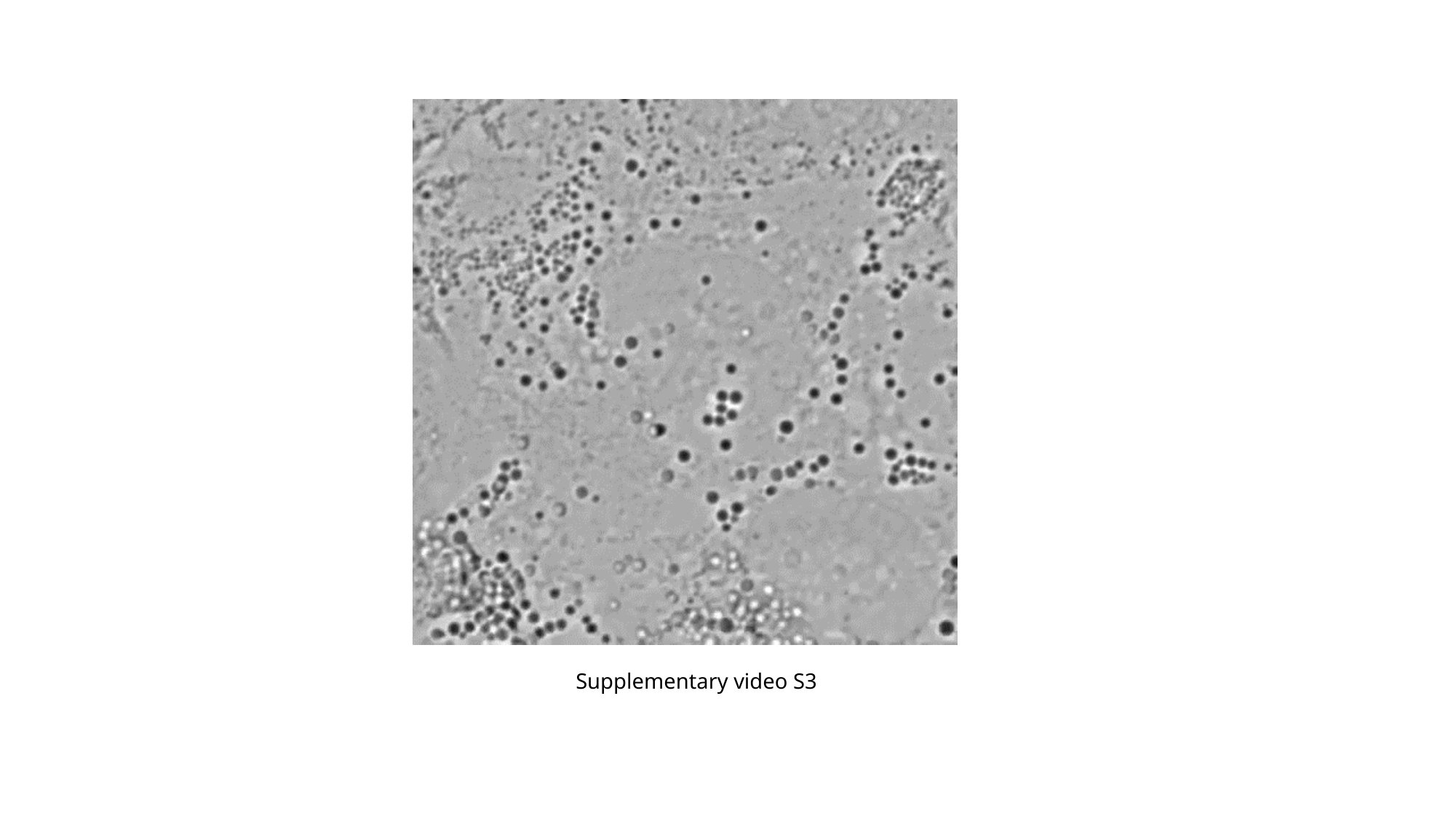

Supplementary video S3

### supplementary video 4

#### Slide 1
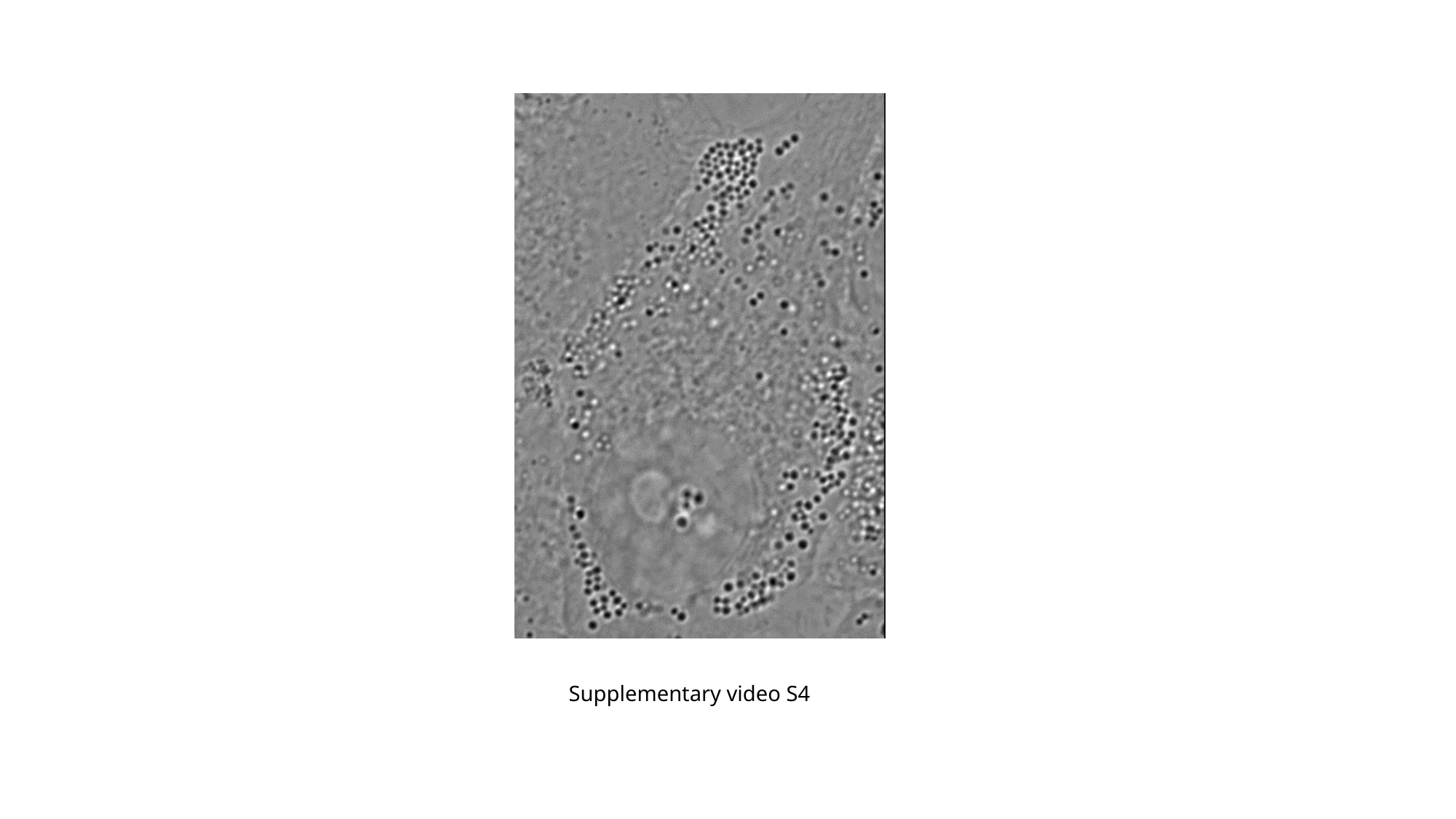

Supplementary video S4

### supplementary video 5

#### Slide 1
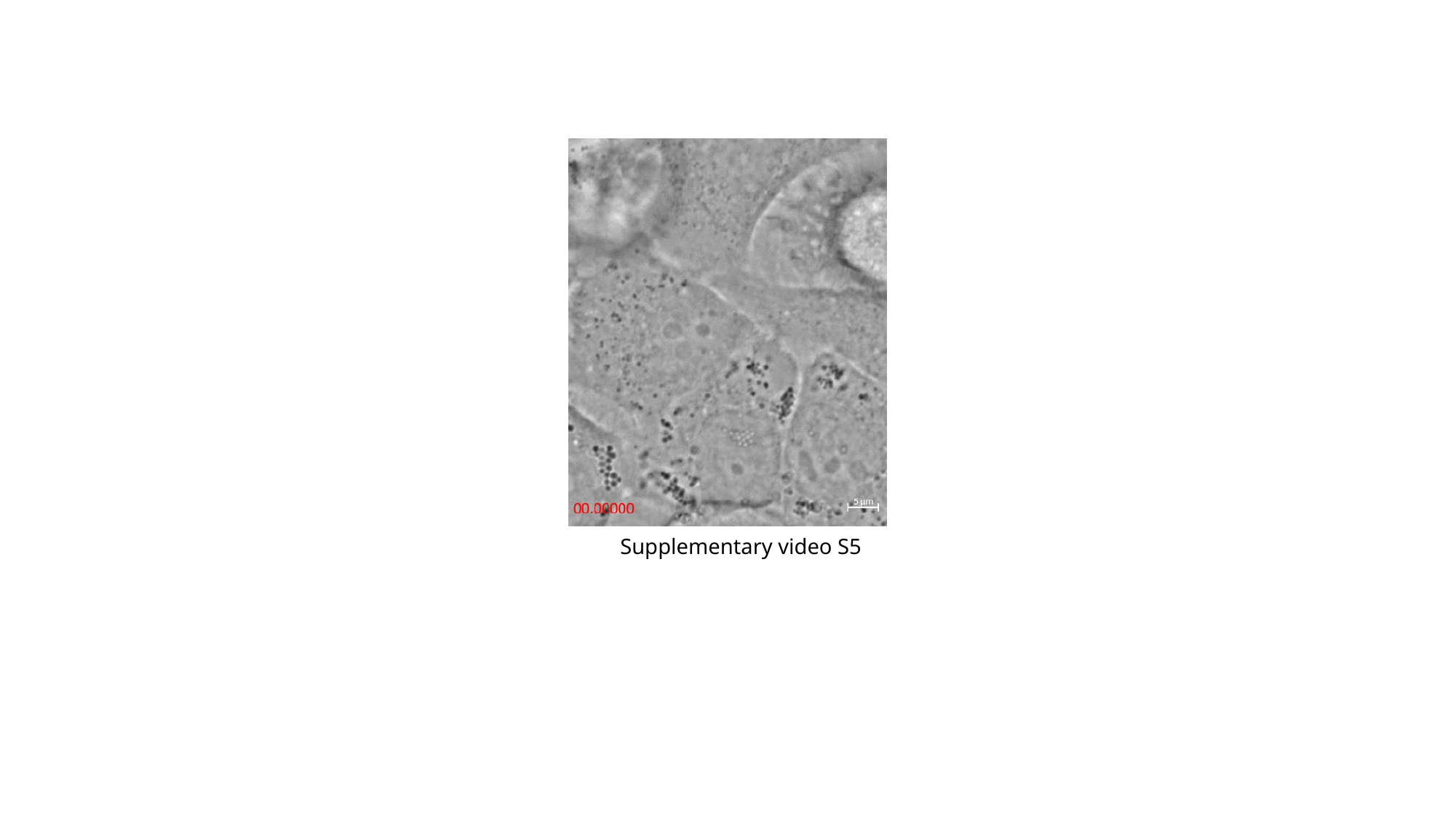

Supplementary video S5

### supplementary video 6

#### Slide 1
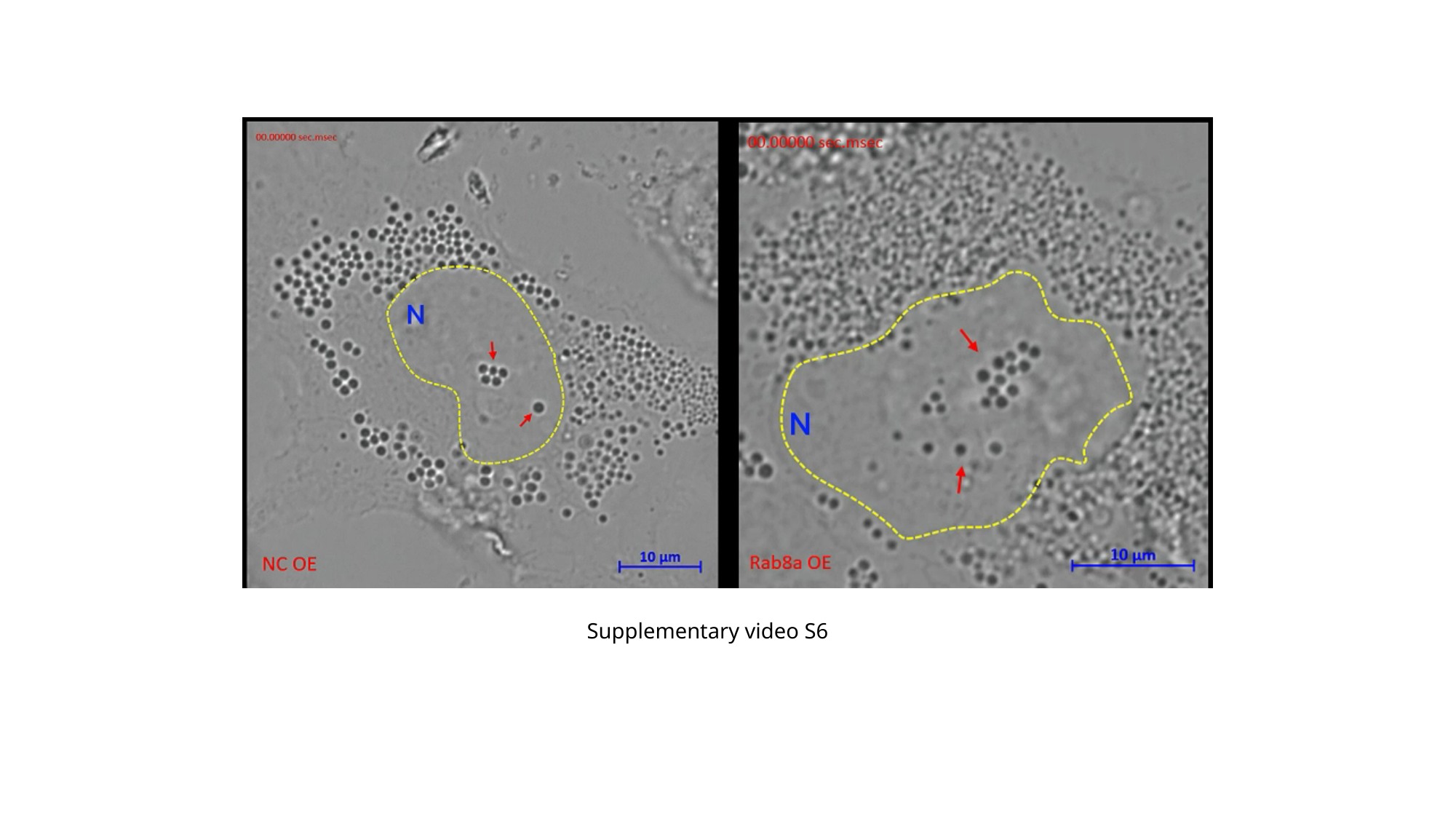

Supplementary video S6
